# A multimodal dataset for reconstructing common marmoset body–environment interactions in a 3D digital-twin framework

**DOI:** 10.64898/2026.06.07.730757

**Authors:** Koki Iwata, Takaaki Kaneko, Catia Correia Caeiro, Diego Thomas, Daisuke Koketsu, Atsushi Nambu, Takako Miyabe-Nishiwaki, Junichi Hata, Ken Nakae

## Abstract

The common marmoset (*Callithrix jacchus*) is an important non-human primate model in neuroscience and biomedical research. However, existing 3D resources for this species have mainly focused on brain atlases or keypoint-based pose estimation, and reusable data resources that jointly describe the body surface, fur, articulated structure, and experimental environment remain limited. Here, we present a multimodal dataset designed to reconstruct body– environment interactions of common marmosets in three dimensions. The dataset includes a whole-body surface mesh derived from computed tomography (CT) images, fur representations based on photographic references, a rigged 3D model for pose-driven animation, synchronized behavioral videos from three individuals recorded for approximately 90 hours from eight viewpoints, 2D and 3D keypoint estimation data, 3D models of the experimental environment constructed from blueprint information, and rendered pseudo-egocentric views generated by integrating pose estimation results with the 3D body and environment models. Technical validation assessed the geometric agreement between the CT-derived mesh and the surface model, the accuracy of 2D and 3D keypoint estimation, the dimensional accuracy of the environment model, and the structural similarity between real and rendered images. This dataset provides a foundation for treating marmoset natural behavior not only as point trajectories but also as a three-dimensional phenomenon involving body shape and its spatial relationship with the environment, thereby enabling applications in behavioral analysis, visualization, synthetic-data generation, and future digital-twin studies.

## Background & Summary

The common marmoset (*Callithrix jacchus*) is small, relatively easy to house and breed, has a short generation time, and is compatible with genetic engineering technologies[1]. At the same time, as a primate it exhibits advanced sensory, cognitive, and social behaviors[2, 3]. For these reasons, the common marmoset has recently become established as an important non-human primate model in neuroscience and biomedical research. In particular, behavioral features such as cooperative breeding, food sharing, and social interaction make this species attractive for investigating the neural bases of human-like social behavior and social cognition, and its use is also expected to expand in disease-model research, including studies of neurodevelopmental and neurodegenerative disorders[3, 4]. Recent work has increasingly emphasized three-dimensional quantification of natural behavior, including studies that infer social, cognitive, and pathological features from 3D pose time series[5] and systems that track the 3D poses of multiple individuals in real time for closed-loop experiments[6].

Existing 3D resources for the common marmoset have primarily focused on neuroanatomical atlases[7, 8, 9, 10, 11, 12] or on pose estimation frameworks aimed at reconstructing keypoints and trajectories[5, 6, 13]. These resources are essential for providing standard reference spaces for the brain and for high-throughput quantification of natural behavior. However, their main purpose is to obtain coordinates in brain space or spatiotemporal coordinates of body parts. They do not describe, as an integrated resource, the external body surface, fur, rig, and the experimental environment with which the animal interacts. Thus, there remains a substantial gap in resources that can represent marmoset behavior not simply as a set of points, but as a visual and spatial phenomenon involving the relationship between the body and the environment.

This gap becomes clearer when compared with recent progress in 3D body-model research in other species. In mice, CT-based three-dimensional virtual animals have been constructed and used to generate synthetic training data and ground-truth labels for 2D and 3D pose estimation[14]. In rats, a biomechanically realistic virtual rodent was trained to imitate natural behavior, and its internal representations were shown to predict the structure of neural activity in behaving rats[15]. In macaques, in addition to OpenMonkeyStudio for 3D pose estimation during free behavior[16], parametric 3D face models have been developed for generating controlled visual and social stimuli[17, 18]. These examples demonstrate the value of 3D information for behavioral measurement and of renderable model resources across species.

A digital-twin framework for common marmoset behavior requires a renderable body model, an articulated control structure, and a spatially matched environment that can be synchronized with measured behavioral data. We constructed a reusable resource for the common marmoset that integrates a CT-based whole-body surface model, fur representations based on photographic references, a rig for synchronizing behavior, and a 3D environment model including the experimental cage and surrounding apparatus. For the fur representation, we implemented a high-fidelity version using Hair Curves in Blender[19], and also prepared a lightweight representation that can be more readily used in other rendering environments, such as Unity and Unreal Engine. This design aims to make the resource usable beyond a single software platform. By synchronizing pose estimation data with the 3D body and environment models, the dataset enables body–environment interactions that are difficult to capture from keypoint sequences alone to be visualized in a spatially coherent manner.

Digital twins are generally understood as frameworks that connect digital representations of real-world objects or systems with measured data for understanding, analysis, and prediction[20]. The present resource is positioned as a foundation for such digital-twin research in marmosets (e.g., emulating a cognitive process of behaving marmosets in silico to understand underlying neural mechanisms)[21]. Rendered videos and web-viewable 3D models generated from these resources are also provided on the project page (https://nakaelab.github.io/marmoset-digital-twin/), allowing users to inspect the reconstructed marmoset behavior and the digital-twin environment interactively. As an example of such visualization, we also provide rendered pseudo-egocentric-view videos that simulate the visual scene from the marmoset’s estimated head-centered viewpoint.

## Methods

### Workflow

The workflow for constructing the dataset is shown in Figure 1a. In the horizontal workflow, the orange branch represents the procedure for generating a 3D marmoset model from CT scan images, the green branch represents 2D and 3D keypoint estimation from videos, and the blue branch represents construction of a 3D environment model from blueprint drawings. Finally, the 3D marmoset model and the 3D environment model were combined to create a 3D digital-twin marmoset that reproduces marmoset behavior in three dimensions, and each component was validated.

**Figure 1.**
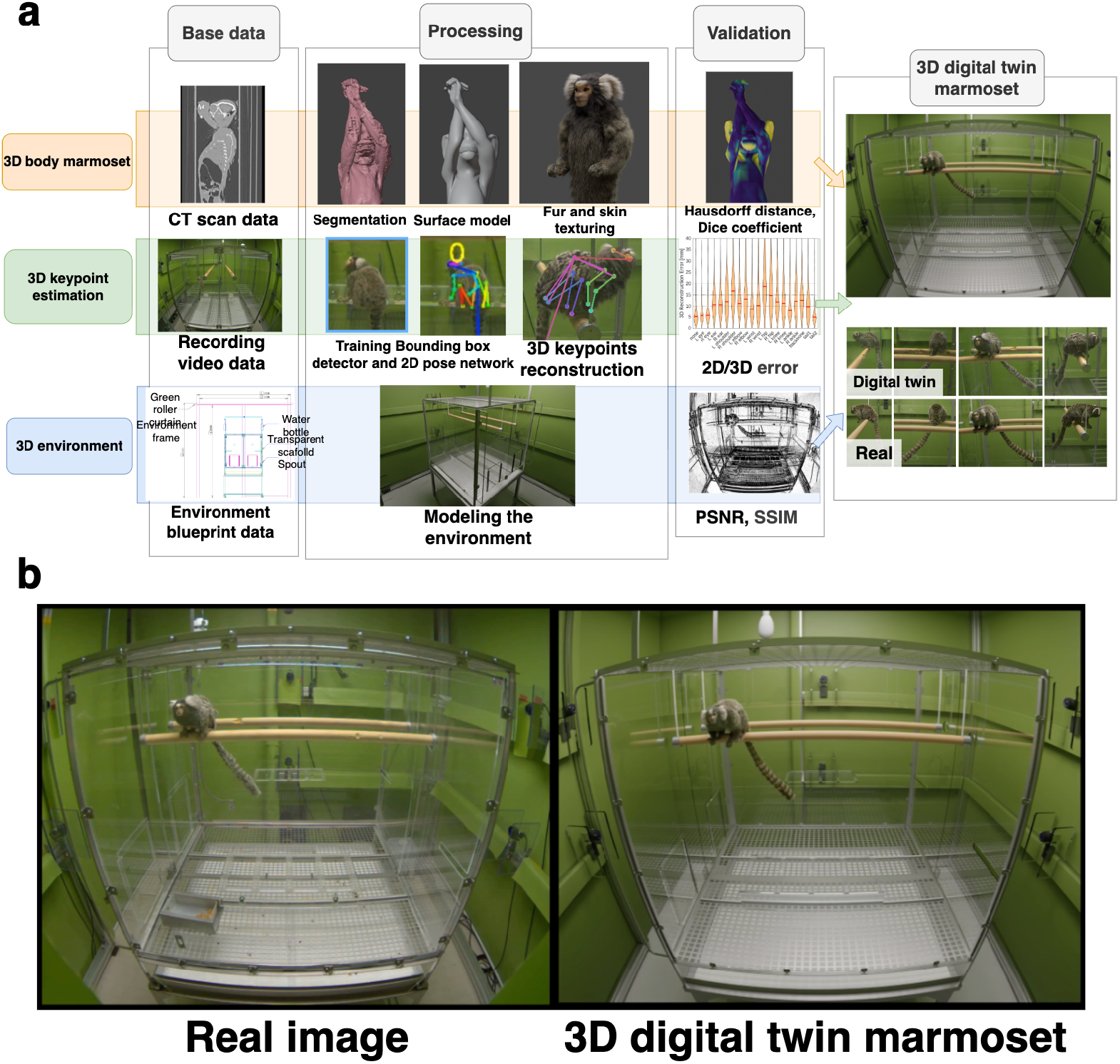
(a) Workflow for constructing the 3D digital-twin marmoset individual and environment, and for creating the dataset. The orange branch shows the method used to create a computer-operable 3D marmoset from CT scan images of the marmoset; the green branch shows the method used to perform 3D pose estimation from videos of the marmoset; and the blue branch shows the method used to create a 3D environment from blueprints of the cage and frame used in the recording environment. For elements not specified in the drawings, approximate positions were determined from images of the experimental environment and finalized together with adjustment of the camera parameters. Some equipment installed in the recording environment, including cameras, was also reproduced in the 3D environment model to create the digital-twin environment. Finally, the 3D marmoset model and 3D environment model were combined to reproduce marmoset behavior in three dimensions, and each method was validated. (b) Comparison between the digital-twin reconstruction generated using the 3D digital-twin marmoset and the real recording environment. The left panel shows real recorded data, and the right panel shows data reproduced in the computer environment.

Figure 1b shows a comparison between the real recording environment and its reconstruction using the 3D digital-twin marmoset. The left panel shows real recorded data, whereas the right panel shows the data reconstructed in the computer environment. Although small discrepancies remain in shape, color, and object placement, the overall appearance of the marmoset and the structure of the recording environment are reproduced. The workflow used to create this 3D digital-twin marmoset is described in the following sections.

### Animals

Two types of data were acquired from common marmosets (*Callithrix jacchus*) in this study: CT imaging data for creating the 3D marmoset model and behavioral video data for estimating 3D pose. Different individuals were used for these two purposes.

For CT imaging, one male common marmoset was used after euthanasia (5 years and 8 months old; body weight, 291 g). CT imaging was performed at the Central Institute for Experimental Medicine and Life Science (CIEM). This individual was housed and managed at CIEM under an experimental protocol approved by the animal experiment committee of that institute (approval no. 14026A).

For behavioral-video recording, three common marmosets (two females and one male), listed in Table 1, were used. These individuals were housed at the Institute for the Evolutionary Origins of Human Behavior, Kyoto University (EHUB). Behavioral recordings were conducted in a dedicated recording space. All animal experiments related to behavioral recording were approved by the Animal Welfare and Animal

**Table 1.** Details of the individual common marmosets recorded in the measurement environment.

| ID | Sex | Date of Birth | Measurement Duration | Measurement Hours |
| --- | --- | --- | --- | --- |
| A | Female | 2018/05/28 | 2021/04/26 – 2021/10/08 | 55.46 |
| B | Male | 2014/04/11 | 2021/08/27 – 2023/03/13 | 33.38 |
| C | Female | 2011/11/21 | 2021/10/09 – 2021/10/10 | 1.33 |

Care Committee of the EHUB (approval nos. 2021-023 and 2022-101). All procedures were conducted in accordance with the Guidelines for Care and Use of Nonhuman Primates established by the center.

### CT image acquisition

Whole-body CT images were acquired using an Aquilion ONE TSX-306A system (Canon Medical Systems). The marmoset was scanned in the posture shown in Figure 1b. The imaging parameters were as follows: tube voltage, 120 kV; tube current, 150 mA; effective tube current–time product, 92 mAs; scan speed, 0.5 s*/*rot; and imaging time, 4.9 s. The acquired CT images had an in-plane resolution of 200 *µ*m *×* 200 *µ*m and a slice thickness of 300 *µ*m. As indices of radiation exposure, CTDI_vol_ was 11.1 mGy and DLP was 343.1 mGy·cm.

### Segmentation of CT data

To extract the body-surface shape of the marmoset from the acquired CT images, we implemented a preprocessing pipeline in MATLAB (Figure 2a). First, the region inside the metal cylinder containing the marmoset was extracted from the raw CT scan data. Next, hole filling and removal of small regions smaller than 100 voxels were applied to the extracted body region using image-processing procedures. Finally, global smoothing was applied to obtain a mask of the marmoset skin surface derived from CT data. Owing to limitations in the spatial resolution of the CT scan and the segmentation procedure, the resulting surface mesh retained jagged shapes around the face and hands. In addition, because the scanned individual had damage to the skull, a hole was present in the head surface. marmoset model.

**Figure 2.**
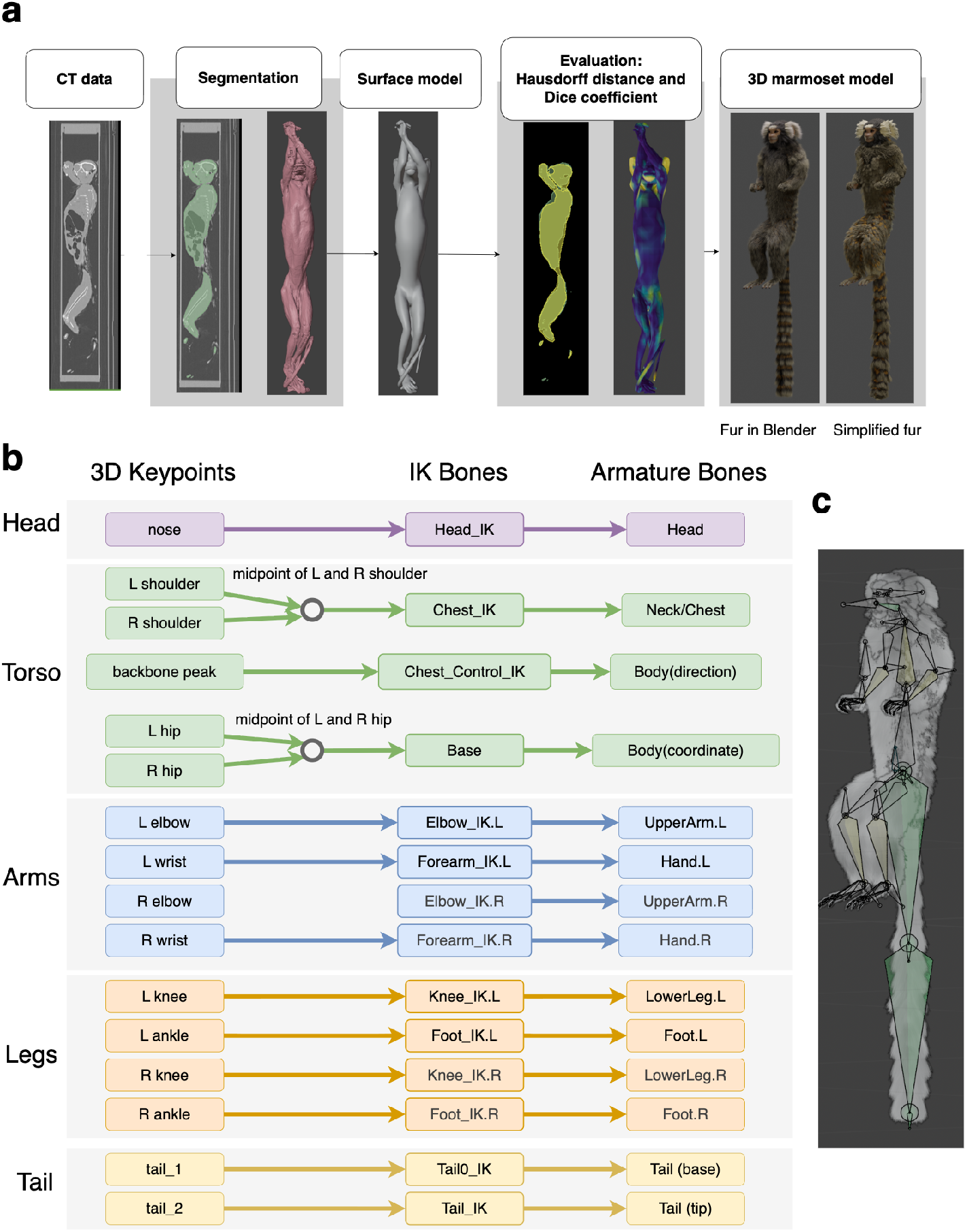
(a) Procedure used to create the 3D marmoset. Segmentation data, shown in green, were generated from CT scan images by image processing. The segmentation was converted into a 3D mesh to obtain the segmentation-derived marmoset surface model shown in red. A manually created marmoset surface model, shown in gray, was then generated to fit the red mesh. A 3D marmoset model was created by applying textured fur and skin to the surface model. A lightweight version of the 3D marmoset model with simplified fur representation was also created. The shape accuracy of the 3D marmoset surface model was evaluated by comparison with the segmentation-derived mesh using the Dice coefficient and Hausdorff distance. (b) Correspondence between the rigged IK bones, armature bones, and 3D keypoints. This panel shows how each bone rigged to the 3D marmoset corresponds to the 3D keypoints described in Figure 3c. Most 3D keypoints and IK bones have a one-to-one correspondence, whereas the shoulder and hip IK bones correspond to the midpoint calculated in 3D space. (c) Armature structure overlaid on the semi-transparent surface and fur models. The visualization shows that bones are embedded in the head, limbs, tail, and other body regions, enabling articulated pose control of the marmoset model.

### Creation of the surface model

Because the segmentation-derived body-surface mesh contained the rough shapes and defects described above, a corrected 3D marmoset surface model was manually created using Blender (Figure 2a). The shape accuracy of the surface model was evaluated by comparison with the segmentation-derived mesh. The Dice coefficient and Hausdorff distance were used as evaluation metrics.

### Application of fur and skin textures

Fur was added to the surface model to reproduce the external appearance of the marmoset (Figure 2a). Fur was generated using the Hair Curves function in Blender. The shape of the fur was manually adjusted with reference to photographs of real marmosets. Fur colors were extracted from photographs of real marmosets using the color-picker function of the GNU Image Manipulation Program and were ultimately determined subjectively. The selected fur colors were applied to Hair Curves in Blender and fine-tuned to match the real appearance. Details of the fur parameters are provided in the publicly available Blender files. In addition to this Hair Curves– based representation, we created a lightweight version of the 3D marmoset model in which the fur was approximated by a general-purpose mesh rather than the Hair Curves function. This simplified representation reproduces the overall fur color and silhouette while remaining usable in rendering environments that do not support Blender’s Hair Curves, such as Unity and Unreal Engine.

### Rigging

Rigging is the process of adding an internal skeletal structure to a 3D surface model so that it can be posed and animated. In this study, rigging was performed to enable the digital-twin marmoset to reproduce marmoset-like movements based on 3D pose estimation results. The internal structure was defined as an articulated hierarchy, or armature, consisting of trunk, limb, tail, and inverse-kinematics (IK) control bones. The 3D marmoset surface model was rigged in Blender by defining an articulated hierarchy (armature). The armature consists of a main trunk hierarchy, Base → Body → Chest → Neck → Head; left and right forelimbs, Shoulder → UpperArm → Forearm → Hand, including bones for each finger; left and right hindlimbs, Pelvis → UpperLeg → LowerLeg → Foot, including bones for each toe; and the tail. In addition, 13 IK bones (Chest_IK, Head_IK, Foot_IK.L/R, Knee_IK.L/R, Forearm_IK.L/R, Elbow_IK.L/R, Tail_IK, Tail0_IK, and Chest_Control_IK) were defined to control animation based on pose estimation results. A complete list of the bone hierarchy is provided in the public repository. The keypoint positions obtained from 3D pose estimation were mapped to the IK-bone positions of the rigged 3D marmoset (Figure 2b, c). This mapping enabled animation of the 3D marmoset model based on pose estimation results.

### Video-data acquisition

To record marmoset behavior simultaneously from eight directions, eight Basler acA2040-35gc cameras (Basler AG) were arranged around the cage (Figure 3a). Each camera was equipped with a Theia ML410 4–10 mm lens. Videos were recorded using the Motif system (Loopbio, Lange G, Wien, Austria). The resolution was set to 2048 *×* 1536 pixels, and the frame rate was set to 24 fps. Although the video data were acquired at 24 fps, they were saved at 25 fps because of the Motif recording system. Therefore, the videos were converted to 24 fps during preprocessing. As shown in Table 1, a total of 222 sessions and 7,790,723 frames of behavioral data (approximately 90 hours) were acquired from the three individuals. By individual, these consisted of 167 sessions from individual A (4,791,838 frames), 51 sessions from individual B (2,883,861 frames), and 4 sessions from individual C (115,024 frames). Camera calibration was performed using the OpenCV framework. Intrinsic parameters, including lens-distortion coefficients, were obtained by inputting images of a checkerboard pattern acquired by each camera into `cv2.omnidir.calibrate`. Extrinsic parameters, including camera positions, were initialized by inputting 3D coordinates of landmark positions in the recording space and their 2D projected coordinates in each camera image into `cv2.solvePnP`. The intrinsic and extrinsic parameters were then jointly optimized by minimizing the projection error of the trajectory of a small object (a table-tennis ball) moved inside the cage, thereby improving calibration accuracy.

**Figure 3.**
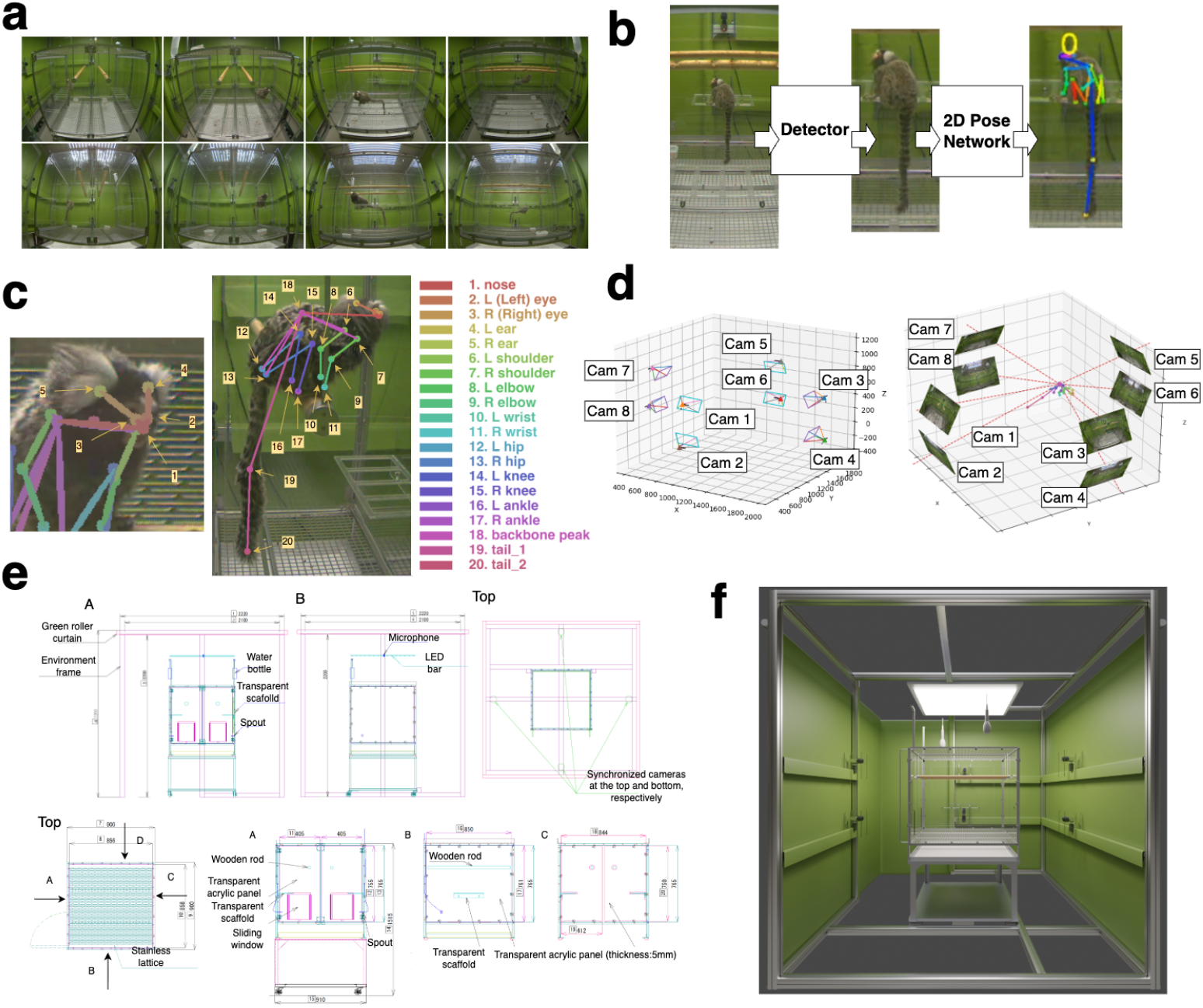
Acquisition of 3D keypoints, cage and frame blueprints. (a) Images of a marmoset recorded from eight camera directions. (b) Method for 2D pose estimation using a bounding-box detector and a 2D pose estimation network applied to the recorded marmoset images. (c) Keypoint body parts defined for pose estimation, their corresponding names and numbers, and the skeleton. The left image shows an enlarged view around the head, and the right image shows the whole body. (d) Method for 3D pose estimation. The left image shows camera positions and orientations estimated by calibration of the cameras used for recording. The right image shows an example of 3D pose estimation using the cameras. 3D pose estimation was performed from the 2D keypoint positions in the real images and the spatial relationships among the cameras. (e) Blueprints of the cage and frame. The green roller curtains, used as a plain background for recording, were not specified in the blueprints; their positions were determined from images of the experimental environment and finalized together with adjustment of the camera parameters. (f) Example of the 3D environment model created from the cage and frame blueprints. The image was rendered from the B direction shown in panel e, with parts of the green curtain, frame, and cameras hidden.

### 2D and 3D keypoint estimation

To reproduce the behavior of living marmosets in the digital-twin marmoset, we used existing 3D pose-estimation techniques as an intermediate representation between behavioral videos and the digital-twin model. Behavioral video data were first converted into keypoint-based 3D pose data, and these 3D pose data were then used as control data for the internal rig of the digital-twin marmoset. To estimate the 3D pose of the marmoset from multi-view videos, we used a three-stage approach. First, bounding boxes were detected in each camera image; second, 2D keypoints were estimated; and third, 3D keypoints were reconstructed from the multi-view 2D keypoints (Figure 3a–d).

#### Bounding-box detection

For each camera image, the marmoset region was detected using a bounding-box detector based on the YOLOX architecture[22] (Figure 3b). The detector was trained using MMDetection (v2.26.0). The training data were prepared by modifying the publicly available Zenodo dataset that contains annotation data[23]. As the pretrained detector, we used a detector from a previous study that did not include the tail. Based on the annotation data and pretrained model, the bounding-box detector was fine-tuned to include the tail. Approximately 3,000 images were used for training, with a train/test ratio of 8:2. Training was completed in approximately 1 h using a GPU (NVIDIA A100). All subsequent mentions of GPU-based training refer to the same environment. The trained model from the epoch that achieved the highest mAP on the test data was selected. Here, mAP denotes mean average precision, a standard object-detection metric based on the COCO evaluation protocol[24]. In this metric, a predicted bounding box is considered correct when it sufficiently overlaps with the manually annotated ground-truth box. The degree of overlap is measured by Intersection over Union (IoU), and mAP summarizes detection performance across multiple IoU thresholds. In this study, mAP was computed as the average across IoU thresholds from 0.50 to 0.95 in increments of 0.05.

#### 2D keypoint estimation

For 2D pose estimation, we adopted the HRNet-W48-DEKR architecture[25] and trained the model using MMPose (v0.29.0). As the pretrained pose estimation model, we used a model from a previous study that did not include the tail, and finetuned it as a 20-keypoint pose estimation model by adding two tail keypoints. The annotation data were generated by annotators who were not specialists in marmosets or behavioral analysis. To standardize the annotation procedure, the annotators were first trained using videos of marmosets in which the target anatomical locations had been visually marked. They then manually annotated the corresponding keypoint locations on images of unmarked marmosets. The number of images used for training was the same as that used for bounding-box training.

Training was completed in approximately 2 hours using the GPU. The trained model from the epoch that achieved the highest mAP on the test data was selected. The keypoint definition used for pose estimation consisted of 20 body-surface landmarks on the marmoset (Figure 3c). The keypoints were nose, L/R eye, L/R ear, L/R shoulder, L/R elbow, L/R wrist, L/R hip, L/R knee, L/R ankle, backbone peak, tail 1, and tail 2. To evaluate the accuracy of 2D keypoint estimation, the mean pixel error on the test data was calculated.

The accuracy of the pose estimation model was also evaluated using mAP based on the COCO evaluation protocol. In pose estimation, mAP is computed using Object Keypoint Similarity (OKS) as the similarity measure instead of IoU, which is used in object detection. OKS is defined as follows:

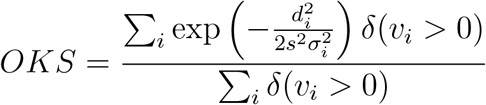

where *d*_*i*_ is the Euclidean distance between predicted keypoint *i* and the groundtruth keypoint, *s* is the object scale (the square root of the segmentation area), *σ*_*i*_ is a keypoint-specific constant, and *v*_*i*_ is the visibility flag of the ground-truth keypoint. OKS ranges from 0 to 1, with values closer to 1 indicating predictions closer to the ground truth. AP was computed from the area under the precision– recall curve at each OKS threshold. In this study, the pose estimation model that achieved the highest mAP was selected. For the values of *σ*_*i*_, we directly adopted the values calculated from inter-annotator standard deviations in the COCO human-pose estimation benchmark for the corresponding keypoints (nose, eyes, ears, shoulders, elbows, wrists, hips, knees, and ankles; Lin et al., 2014). For the additional keypoints (back, tail_1, and tail_2), we set *σ* = 0.089. These *σ* values were not calculated from marmoset-specific annotation variance, and the difficulty of localizing keypoints may differ between marmosets and humans.

#### 3D keypoint reconstruction

Finally, 3D keypoint coordinates were reconstructed using the 2D keypoints estimated from the eight cameras and the camera parameters obtained by calibration (Figure 3d). 3D reconstruction was performed using the Zenodo resource that contains the execution scripts[26]. In brief, for each frame, bounding boxes across different cameras were grouped based on geometric consistency, and 3D poses were constructed by triangulation. Next, 3D poses between adjacent keyframes were temporally associated using the Hungarian algorithm. Individual IDs were then assigned to each 3D tracklet, fragmented tracklets were merged, and final smoothing was performed. The 3D pose time-series data for each individual were ultimately smoothed and normalized spatiotemporally using Anipose[27]. Detailed algorithms and hyperparameters for each step are described in Kaneko et al.[5].

To evaluate the accuracy of 3D keypoint reconstruction, we calculated the error between 3D keypoints reconstructed from manually specified 2D keypoints, which served as the ground truth, and 3D keypoints reconstructed from 2D keypoints estimated by the 2D keypoint estimation model. In addition, to evaluate the quality of the estimated 3D keypoints in actual data, we calculated the reprojection error. Reprojection uses the camera-calibration data, including camera positions in 3D space and lens distortion, to correct image distortion and transform 3D coordinates into 2D camera coordinates. The reprojection error was calculated as the difference between the coordinates obtained by reprojecting the 3D keypoints into 2D keypoints and the coordinates of the estimated 2D keypoints.

### Construction of the 3D environment

Blueprint information for the cage and the outer frame—a structural framework surrounding the recording area, on which the cameras were mounted—was used to construct the 3D environment model (Figure 3e). Numbers were assigned to the parts in the blueprints, and dimensional information for the cage and outer frame was extracted. Modeling was performed using Blender v4.5.0, and 3D models of the cage and frame were created based on the extracted dimensions. The dimensions of each part of the created model were measured using the Python functionality built into Blender and a custom Python program, and the model was adjusted to match the blueprint values. Because the blueprints did not include the green curtains in the background, the green sheets near the cameras, or the height of the frame on which the cameras were installed, these elements were created to match the real recording images as closely as possible. The cage was placed within this environment, and the initial placement was determined so that the rendered views approximately matched the images from the recording cameras.

The camera-distortion parameters and the 3D camera positions were then manually adjusted so that the real recorded images and rendered images were visually consistent. The colors of the curtains and lighting in the experimental environment were also adjusted by comparing real recorded images with rendered images and tuning the material colors and light intensity in Blender. This procedure produced the 3D environment model shown in Figure 3f. The figure shows the 3D environment model from the B direction indicated in Figure 3e, with parts of the green curtains and frame hidden. The final 3D environment, measurement scripts, and camera parameters are provided in the public repository.

### Construction of the 3D digital twin

The 3D marmoset model, 3D keypoints, and 3D environment model were combined to construct a 3D digital-twin environment for reproducing marmoset behavior in three dimensions (Figure 1b). As a fine-adjustment step, the rigged 3D marmoset model was controlled by its IK bones based on the positions of the 3D keypoints, thereby reproducing the marmoset’s behavior in 3D. Because the 3D keypoints were defined in absolute coordinates in the real space whereas the IK bones of the 3D marmoset model in Blender were controlled in relative coordinates with respect to the base bone of the model, the 3D keypoint coordinates were transformed into the coordinate system of the marmoset model in Blender before applying the keypoints. As a result, the 3D marmoset model could move according to the 3D keypoint positions. A digital-twin environment was also constructed in the same manner using the simplified 3D marmoset model, providing a lightweight version that can be reused more readily across different rendering environments. The real video corresponding to Figure 1b is available in the repository. To quantify the similarity between real and rendered video frames, we computed the structural similarity index measure (SSIM) and peak signal-to-noise ratio (PSNR). SSIM was used as a perceptual measure of structural similarity between images, following Wang et al. [30], whereas PSNR was used as a conventional full-reference pixel-level fidelity metric [31].

### Pseudo-egocentric view estimation

To estimate the viewing direction of the marmoset, a pseudo-egocentric view vector was generated from the estimated positions of the ears and eyes (Figure 4a). The pseudo-egocentric view direction was defined as the vector from the midpoint of the two ears to the midpoint of the two eyes, and this direction was used to determine the orientation of the marmoset-view camera. The camera origin was set to the estimated midpoint of the two eyes. Because this direction does not necessarily correspond to the actual gaze direction of the marmoset, it should be interpreted as a pseudo-egocentric view estimate.

**Figure 4.**
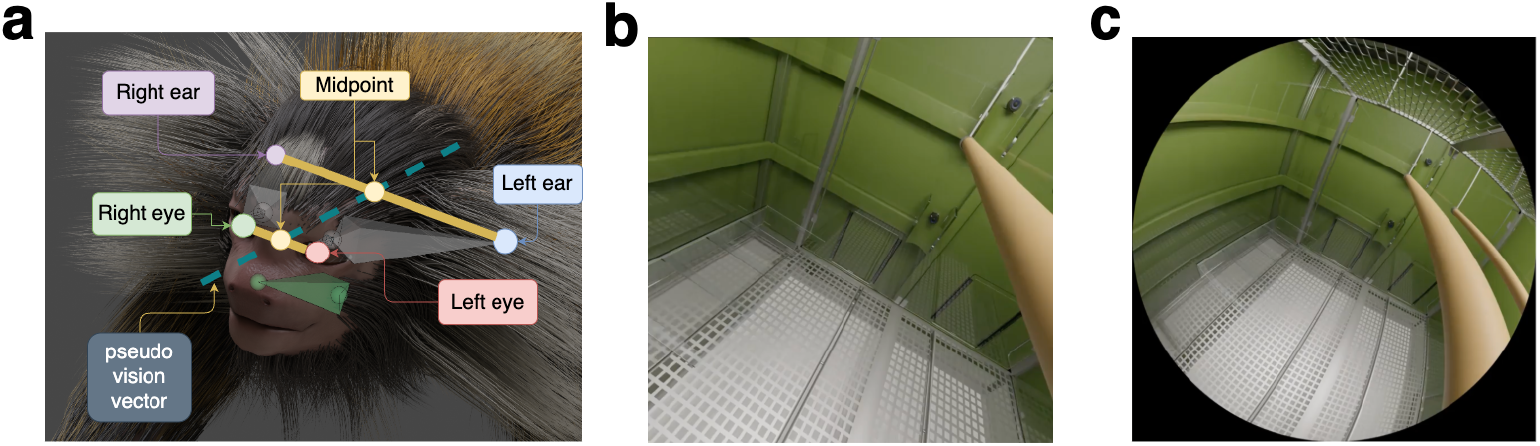
A method for estimating the simulated viewpoint of a 3D digital-twin marmoset and rendering pseudo-egocentric viewpoint images. (a) The pseudo-egocentric view vector was defined as the vector from the midpoint of the ears to the midpoint of the eyes in the 3D marmoset model. (b) Rendered pseudo-egocentric viewpoint image generated using an equidistant camera in Blender. (c) Rendered pseudo-egocentric viewpoint image generated using a pinhole camera in Blender.

According to Troilo et al.[28], the monocular visual field of the common marmoset is approximately − 60° to 100° horizontally and − 70° to 70° vertically. They also reported a retinal area of 209.5 mm^2^ and a posterior focal length of 10.03 mm. In this study, we generated two types of pseudo-egocentric viewpoint renderings using Blender camera models: an equidistant camera and a pinhole camera. The pinhole camera corresponds to the Perspective camera in Blender. For the equidistant camera, the field of view was set to − 70° to 70° in both the horizontal and vertical directions (Figure 4b). For the pinhole camera, the sensor size and focal length were configured based on the reported retinal area and posterior focal length (Figure 4c). Rendering was performed on a GPU using Blender’s Cycles rendering engine. The image resolution was set to 1000 *×* 1000 pixels, and the maximum number of samples was set to 500. The rendered images were exported as an MP4 video at 24 fps using FFmpeg with the H.264 codec and YUV 4:2:0 pixel format. In this study, one recording session was rendered.

## Data Records

Part of the 3D digital-twin marmoset dataset is available through Zenodo, and the complete dataset is available through AWS https://digitaltwin-marmoset.s3.ap-northeast-1.amazonaws.com/index.html. The released data include CT data, training data for 2D pose estimation, video data from three individuals, the trained 2D keypoint estimation model, 2D and 3D keypoint time-series data, rendered pseudo-egocentric view data, and data used to calculate SSIM and PSNR. The folder structure of the 3D digital-twin marmoset dataset is shown in Figure 5. The released data consist of four subfolders: 3D_marmoset_body_data, marmoset_keypoint_data, 3D_environmental_data, and 3D_digital_twin_data.

**Figure 5.**
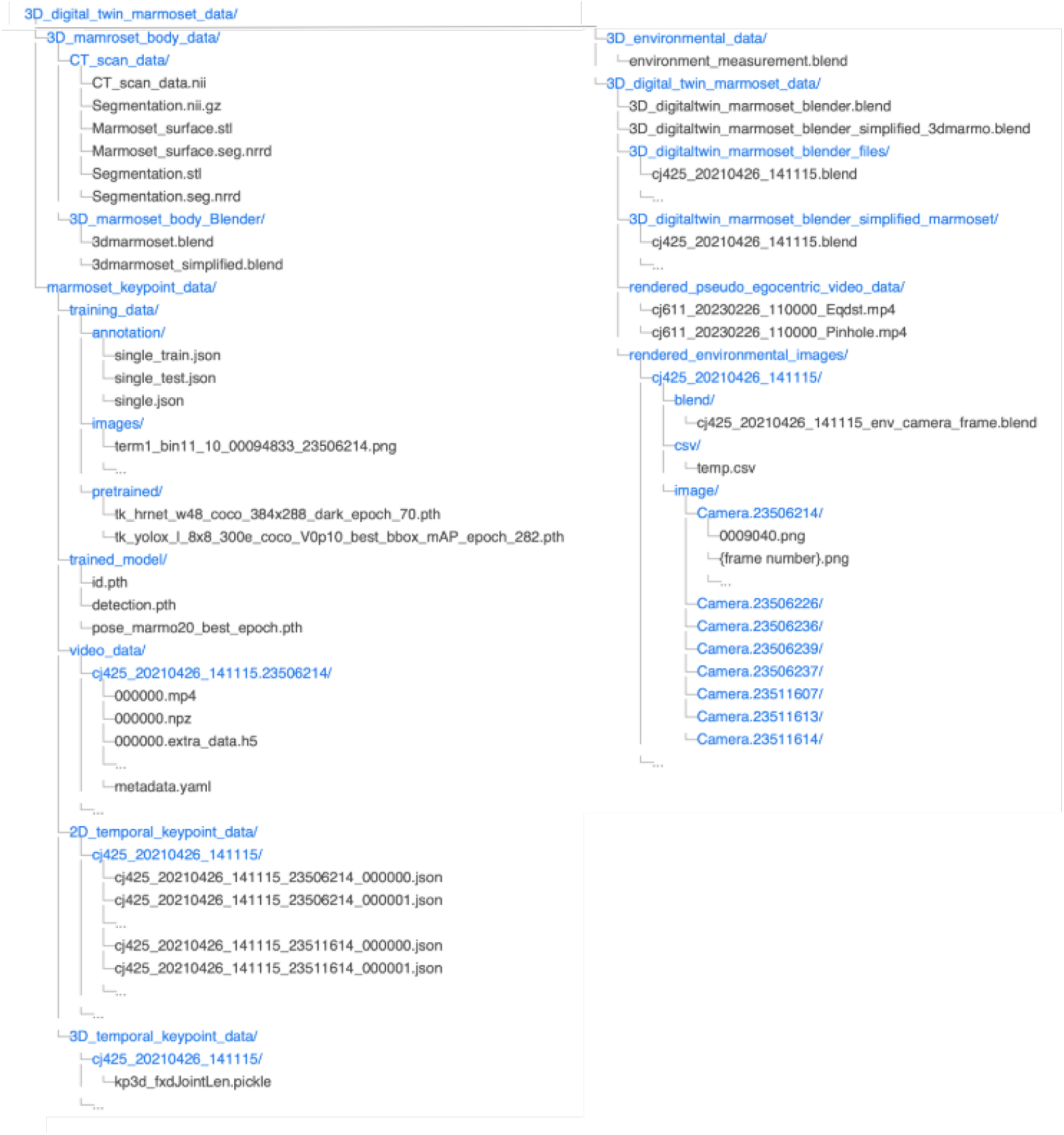
Folder structure of the 3D digital-twin marmoset dataset. The folder consists of four subfolders: 3D_marmoset_body_data, marmoset_keypoint_data, 3D_environmental_data, and 3D_digital_twin_data. The 3D_marmoset_body_data folder contains CT data, segmentation data, and surface-model data. The marmoset_key-point_data folder contains annotation and image data used for training, video data, trained models, and estimated 2D and 3D keypoint data. The 3D_environmental_data folder contains Blender files of the 3D environment model. The 3D_digital_twin_data folder contains Blender files that integrate all components, rendered pseudo-egocentric view data, and data used to calculate SSIM and PSNR.

### 3D marmoset data

#### CT data

CT data are stored in the CT_scan_data folder within 3D_marmoset_body_data. The CT data are stored under the filename CT_scan_data.nii. The post-segmentation data are stored under the filename Segmentation.nii.gz. The segmentation mesh data are stored under the filename Segmentation.stl. The surface-model data are stored under the filename Marmoset_surface.stl. Because the comparison based on the Dice coefficient and Hausdorff distance requires specific segmentation data, both the segmentation data and surface-model data are also stored in the .nrrd file format. The CT data and segmentation data stored in NIfTI format are saved as .nii and .nii.gz files. The segmentation mesh data and surface-model data are saved in STL format.

#### 3D marmoset body

The 3D marmoset body models are stored in the 3D_marmoset_body_Blender folder within 3D_marmoset_body_data and are saved in Blender format (.blend). Each file contains only the 3D marmoset model itself, without the environment model. Two files are provided: 3dmarmoset.blend, which contains the high-fidelity model whose fur was created with the Hair Curves function, and 3dmarmoset_simplified.blend, which contains the lightweight model whose fur was approximated by a simplified mesh.

### Marmoset keypoint data

#### Training data

The training data are stored in the training_data directory within marmoset_keypoint_data. This directory is organized into three subdirectories: annotation, image, and trained_model. The annotation directory contains annotation files used to train and evaluate the bounding-box detector and 2D keypoint estimation model, whereas the image directory contains the corresponding image data. The trained_model directory contains the pretrained detection and pose-estimation models generated by Kaneko et al. [5].

The annotation files are provided in JSON format following the COCO dataset structure. Two files are included: single_train.json and single_test.json, corresponding to the training and test sets, respectively. These files contain annotations for approximately 3,000 images, which were divided into training and test sets at a ratio of 8:2. Each JSON file consists of three main fields: images, annotations, and categories. The images field records the file name, image ID, and image resolution, including width and height. The annotations field stores the bounding-box coordinates in COCO format, [x, y, width, height], together with 20 keypoints for each animal, represented as repeated [x, y, visibility] entries. The categories field defines the marmoset category, keypoint names, and skeleton connectivity.

Image files are stored in PNG format (.png). The pretrained model files are stored in PyTorch format (.pth) and were created in formats used by the MMDetection and MMPose frameworks.

#### Trained models

The trained models are stored in trained_model and include weights (learned parameters) for the trained 2D keypoint estimation model, the ID model, and the bounding-box detector. The weights for the 2D keypoint estimation model, ID model, and bounding-box detector are stored in pth format. The 2D keypoint estimation model is stored as pose_marmo20_best_epoch.pth, the ID model as id.pth, and the bounding-box detector as detection.pth.

#### Video data

The video_data folder contains the behavioral video data. Video data are stored in folders named by combining the individual ID, recording date, recording start time, and camera identifier. The video-data folders are named individual ID_yyyymmdd _hhmmss.camid. The individual IDs are A, B, and C in Table 1; the corresponding original IDs are cj611, cj425, and cj357, respectively. Here, yyyymmdd denotes the recording date, hhmmss denotes the recording start time, and camid denotes the camera identifier used in the folder names. Videos recorded from a given start time, such as 090000, are stored in the corresponding folder and split into chunks of 10,000 frames.

The video files are stored in MPEG-4 format (.mp4) using the AVC/H.264 video codec. Each video has a resolution of 2048 *×* 1536 pixels with a 4:3 aspect ratio and a frame rate of 25 fps. The videos are stored as progressive-scan, 8-bit YUV 4:2:0 video streams. The dataset includes recordings from eight cameras. In Figure 3, these cameras are labeled as cam1 to cam8 for readability; however, these labels are not identical to the camid values used in the folder names. The correspondence between the figure labels and the actual camid values is provided in Table 2.

**Table 2.** Camera numbering correspondence.

| Camera Number | Camera ID |
| --- | --- |
| cam1 | 23506226 |
| cam2 | 23511607 |
| cam3 | 23506214 |
| cam4 | 23506236 |
| cam5 | 23511614 |
| cam6 | 23506237 |
| cam7 | 23511613 |
| cam8 | 23506239 |

Each parent folder contains the recorded video data (.mp4) and Motif-system log data (.npz and .h5). The recorded video data are split into chunks of 10,000 frames; when a recording ended before reaching 10,000 frames, the file contains the corresponding number of frames. The video data have a resolution of 2048 *×* 1536 pixels, a frame rate of 25 fps, and H.264 encoding. Each npz file corresponds to a video chunk (for example, 000000.npz corresponds to 000000.mp4) and contains metadata such as frame-by-frame timestamps and camera exposure times within the video chunk. Each h5 file also corresponds to a video chunk (for example, 000000.extra data.h5 corresponds to 000000.mp4) and contains information on frame-by-frame camera-acquisition timing within the video chunk.

#### 2D keypoint data

The 2D_temporal_keypoint folder contains the estimated 2D keypoint-coordinate data. The actual 2D keypoint data in JSON format are stored in folders named {individual ID}-{yyyymmdd}-{hhmmss}. The JSON files are named {individual ID}-{yyyymmdd}-{hhmmss}-{camid} .json. The 2D keypoint data were output by 2D keypoint estimation from the video data, and the parent-folder name of the video data was used as the filename. Each JSON file contains the coordinates of 20 keypoints estimated for each frame of the video data.

#### 3D keypoint data

The 3D keypoint data are stored in the 3D_temporal keypoint directory. The data are organized into folders named individual ID_yyyymmdd_hhmmss, following the same naming convention as the corresponding video data. Each folder contains a single pickle file that stores the 3D keypoint data estimated for each frame of the corresponding video.

Each pickle file contains a dictionary with four keys: kp3d, kp3d_score, kp3d_err, and joint_len. The kp3d field stores the estimated 3D coordinates of the 20 keypoints. The kp3d_score field stores the confidence score for each estimated keypoint, and kp3d_err stores the reprojection error. The joint len field stores the sigma values used for OKS calculation.

### 3D environment data

The environment_model_Blender folder contains the environment_measurement.blend file. This Blender file contains a 3D model of the cage created from blueprint information, as well as a 3D model of the recording space. The file is stored in Blender format (.blend). In the Blender file, the cage model and the recording-space model are organized into separate collections using Blender’s collection function. The file includes only objects defined in the blueprint information and does not include objects not specified in the blueprints, such as lights and cameras.

### 3D digital twin data

The 3D_digitaltwin_marmoset Blender folder contains two base files that integrate the 3D marmoset model with the 3D environment model. The file 3D_digitaltwin_marmoset_blender.blend uses the high-fidelity marmoset model whose fur was created with the Hair Curves function, whereas 3D_digitaltwin_marmoset blender_simplified_3dmarmo.blend uses the lightweight marmoset model whose fur was approximated by a simplified mesh. Session-specific .blend files were created from these base files by applying the estimated 3D keypoints to the 3D marmoset model, thereby representing the reconstructed marmoset motion for each recording session. The session files based on the high-fidelity model are stored in the 3D_digitaltwin_marmoset_blender_files folder, and those based on the simplified model are stored in the 3D_digitaltwin_marmoset_blender_simplified_marmoset folder.

#### Rendered pseudo-egocentric view data

Rendered pseudo-egocentric view data are stored in rendered_pseudo_egocentric_video_data. The video files, cj611_20230226_110000_Eqdst.mp4 and cj611_20230226_110000_Pinhole.mp4, correspond to the two pseudo-egocentric viewpoint renderings described in the Methods section: the equidistant camera and the pinhole camera, respectively. The pinhole camera corresponds to the Perspective camera in Blender. The videos are stored in MP4 (H.264) format at a frame rate of 24 fps, with the pixel format set to YUV 4:2:0.

#### Rendered 3D environment images

The rendered environmental images folder contains rendered image data and the associated files used to calculate SSIM and PSNR. The data are organized by recording session, with each session stored in a folder named according to the recording information, such as cj425_20210426_141115. Each session folder contains three subfolders: blend, csv, and image. The blend folder contains the Blender file used to render the images, such as cj425_20210426_141115_env_camera_frame.blend. The csv folder contains the CSV file storing the 3D pose data applied to the marmoset model in Blender. The image folder contains rendered PNG images, which are further organized into camera-specific folders named by camera identifiers, such as Camera.23506214. Each camera folder contains rendered images for individual frames, named by frame number, such as 0036082.png or frame number.png.

## Technical Validation

### 3D marmoset

The 3D marmoset was manually created based on the CT scan, and the skin-surface mesh from the CT scan was compared with the skin-surface mesh of the created 3D marmoset. The Dice coefficient and Hausdorff distance were calculated using the SlicerRT toolkit[29], an extension for 3D Slicer. The Dice coefficient indicates the degree of overlap between two regions, with values closer to 1 indicating greater overlap. The Hausdorff distance indicates the distance between boundary points of two regions, with values closer to 0 indicating that the two regions are closer.

Figure 6a shows a visualization of the Hausdorff distance between the CT-scan marmoset and the 3D marmoset surface model created in this study. A heat map was generated by comparing the CT-scan marmoset, shown as the red mesh, with the 3D marmoset surface model. The heat map was rendered on the 3D marmoset surface model. The 95th percentile of the Hausdorff distance was approximately 3.41 mm. The heat map represents the Hausdorff distance, and the upper limit was set to 3.5 mm, near the 95th percentile. Yellow regions showing relatively large deviations from the CT scan were located around the ears, back, tail base, and jaw. For the ears, the CT scan did not clearly depict the ear regions, so the ears were shaped based on photographs of real marmosets. For the back, the segmentation after CT-data processing was asymmetric, and the surface model was modified to be bilaterally symmetric during model creation. The evaluation metrics were a Dice coefficient of 0.891 and a mean Hausdorff distance of 1.14 mm (95th percentile, 3.42 mm), indicating a high volumetric agreement between the CT-derived mesh and the manually created mesh. However, the maximum Hausdorff distance was 15.35 mm, indicating local shape discrepancies. These discrepancies were mainly attributable to noise from segmentation around the head. Additional discrepancies were observed around the feet and trunk and around the hands and feet. The former are attributable to noise from segmentation processing, whereas the latter reflect limitations in the modeling procedure.

**Figure 6.**
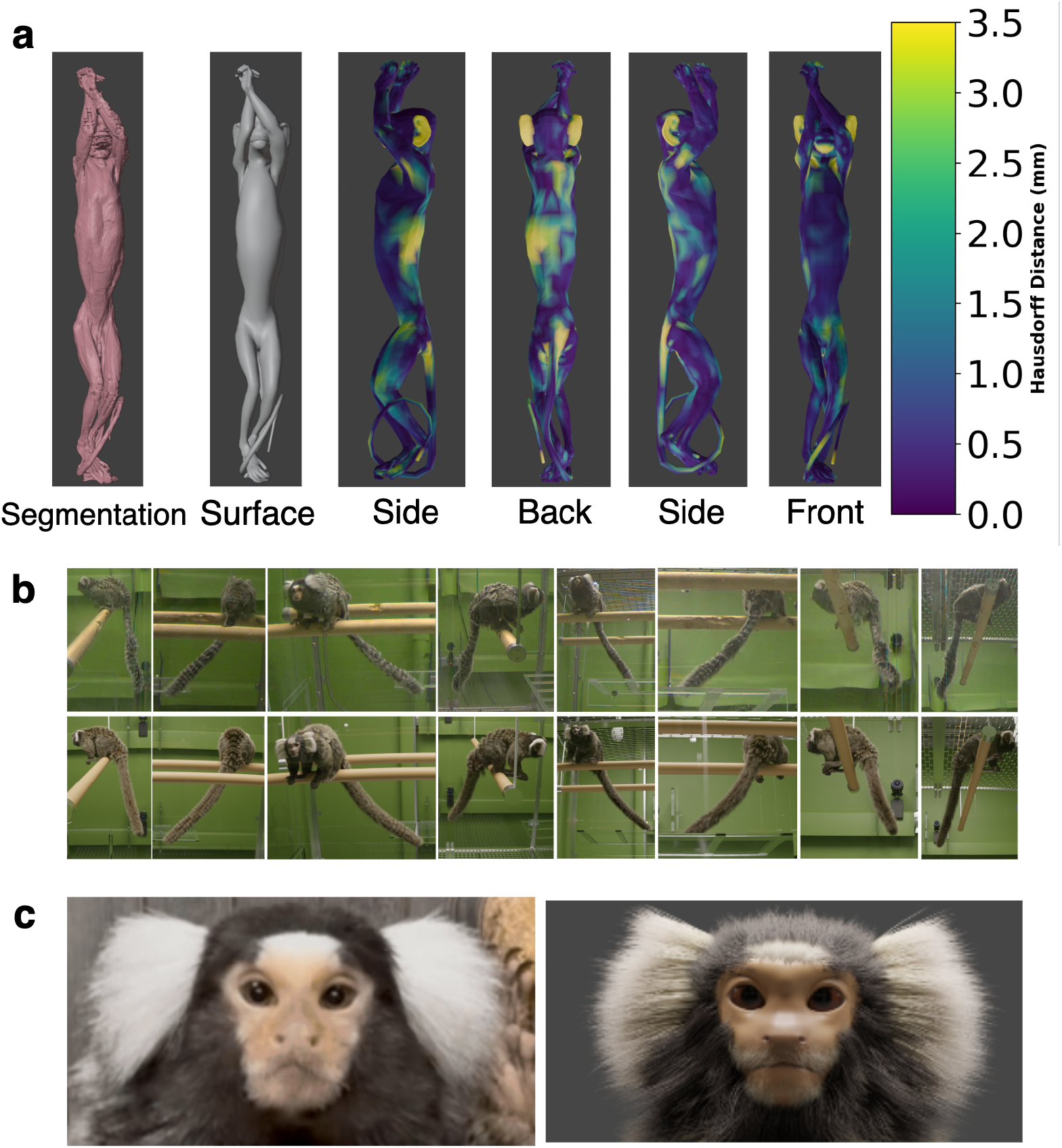
Visualization of the Hausdorff distance and comparison of the appearance of the real marmoset and the 3D marmoset. (a) Visualization of the Hausdorff distance between the CT scan and the 3D marmoset surface model created in this study. The surface model obtained by segmenting the marmoset CT scan and the 3D marmoset surface model are shown on the left. The heat map is drawn on the 3D surface model and represents the Hausdorff distance. The upper limit of the heat map was set to 3.5 mm, near the 95th percentile. (b) Comparison between an image of a real marmoset and an image of the 3D marmoset. The upper image shows the real marmoset, and the lower image shows the 3D marmoset in which the skin was created from the CT scan and fur was added. (c) Comparison between close-up images of the face of the real marmoset and the face of the 3D marmoset. The left image shows the face of the real marmoset, and the right image shows the face of the 3D marmoset.

Figures 6b and c show visual comparisons between a real marmoset and the 3D model. The overall body shape and fur appearance are reproduced, whereas the local discrepancies described above were treated as noise and were not reflected in the created surface mesh. In addition, although the evaluation was subjective, the sizes and positions of the eyes, mouth, and ears in the face were modeled to resemble those of the real marmoset as closely as possible.

### Estimated 2D and 3D keypoints

Figure 7a shows the magnitude of the 2D keypoint error on the test data used for training the pose estimation model. The pose estimation model used was the model from the epoch with the highest mAP. The mean error was approximately 10.15 px. By body part, the facial keypoints (nose: 4.40 px; L/R eye: 3.28/3.34 px) showed the lowest errors; the ears (L/R ear: 6.06/6.70 px) showed intermediate errors; and the limbs (shoulder, elbow, wrist, hip, knee, and ankle: 8.56–12.86 px) and trunk/tail (backbone: 14.73 px; tail1/tail2: 16.12/16.70 px) showed larger errors.

**Figure 7.**
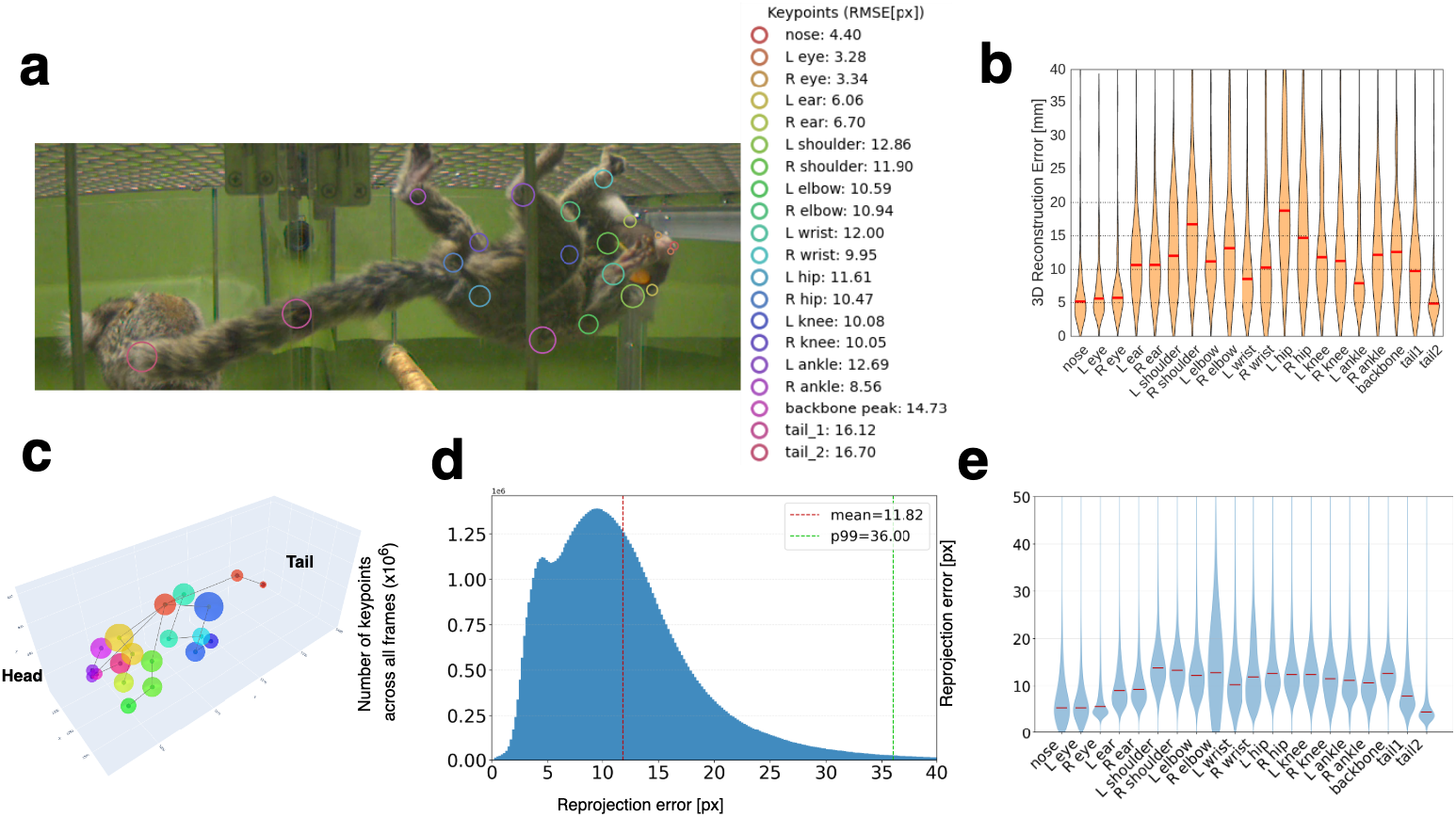
Validation results for the created 2D and 3D keypoints. (a) RMSE [px] against the test data used for training 2D keypoint estimation. Circles on the image are centered on manually annotated keypoints, with each radius set to the mean RMSE for that keypoint. The actual mean errors are shown in the legend. (b) Error [mm] of the 3D keypoints relative to the test data in 3D space. The horizontal axis shows keypoint number, and the vertical axis shows the error in millimeters. The red line shows the mean across keypoints. (c) Visualization of the magnitude of the 3D keypoint error shown in panel b. (d) Distribution of reprojection errors obtained by reprojecting 3D keypoints from all frames into the corresponding camera images. The horizontal axis shows the reprojection error [px], and the vertical axis shows the error frequency. (e) Reprojection error for each marmoset keypoint. The horizontal axis shows keypoint number, and the vertical axis shows the error in pixels. The red line shows the mean error for each keypoint.

Figure 7b shows the magnitude of the mean 3D keypoint error. The error magnitude was calculated from the difference between ground-truth and estimated values. The ground truth was defined as the 3D coordinates triangulated from manually annotated 2D keypoints, whereas the estimated values were defined as the 3D coordinates triangulated from 2D keypoints estimated by the pose estimation model. The difference between these values was defined as the error. The mean 3D keypoint error in this dataset was approximately 12.73 mm (range, 6.89–21.51 mm). This corresponds to approximately 6.4% of the common marmoset body length of approximately 200 mm. By body part, the facial keypoints (nose: 6.89 mm; L/R eye: 7.27/7.55 mm) showed the lowest errors, whereas the limbs and trunk were generally in the range of 10–19 mm. The body-part-specific trends of 2D and 3D errors were generally consistent; for example, facial keypoints with high 2D estimation accuracy also showed small 3D errors, whereas limbs and trunk keypoints with larger 2D errors also showed larger 3D errors. As a reference, Figure 7c shows the 3D keypoint errors visualized in three-dimensional space.

### Reprojection error

#### Camera-calibration reprojection error

As an additional validation of camera calibration accuracy, we evaluated the reprojection error of a marker-derived point moving in the recording space. The 2D marker positions detected in multiple camera views were triangulated into 3D points using the optimized camera parameters, and the reconstructed 3D points were then reprojected back into each camera image. The mean reprojection error was 11.24 px for the in-sample evaluation and 14.57 px for the leave-one-out evaluation, with median errors of 7.27 px and 9.23 px, respectively. These errors correspond to less than 1% of the original 2048 *×* 1536 image size, indicating that the optimized multi-camera calibration was sufficiently accurate for 3D pose reconstruction, although not at a strict pixel-level precision.

#### 3D keypoint reprojection error

The 3D keypoint errors on the test data described above were evaluated only on a limited number of annotated frames. To assess the quality of the full dataset, we calculated the reprojection error across all frames. Reprojection error was measured using the camera-calibration data and the 2D and 3D keypoint data estimated for all frames. The distribution of reprojection errors across all cameras is plotted in Figure 7d. The mean reprojection error was approximately 11.82 px, corresponding to approximately 0.6% of the horizontal image resolution of 2048 px. The 95th percentile was approximately 36.00 px or less. These results indicate that stable 3D reconstruction was achieved for the majority of frames.

**Table 3.** Blueprint vs. 3D model size comparison.

| Blueprint object No. | MarmoCage<br>blueprint<br>[mm] | 3D model<br>size [mm] | Absolute dif-<br>ference [mm] | Absolute ratio<br>(3D/blueprint) |
| --- | --- | --- | --- | --- |
| 1 | 2220 | 2220.00 | $< 10^{-3}$ | 1.00 |
| 2 | 2100 | 2111.11 | 11.11 | 1.01 |
| 3 | 2200 | 2205.28 | 5.279 | 1.00 |
| 4 | 2260 | 2260.00 | $< 10^{-3}$ | 1.00 |
| 5 | 2220 | 2220.00 | $< 10^{-3}$ | 1.00 |
| 6 | 2100 | 2111.55 | 11.55 | 1.01 |
| 7 | 900 | 900.00 | $< 10^{-3}$ | 1.00 |
| 8 | 856 | 871.07 | 15.07 | 1.02 |
| 9 | 900 | 900.00 | $< 10^{-3}$ | 1.00 |
| 10 | 858 | 874.95 | 18.94 | 1.02 |
| 11 | 405 | 420.74 | 15.74 | 1.04 |
| 12 | 755 | 755.00 | $< 10^{-3}$ | 1.00 |
| 14 | 1515 | 1515.00 | $< 10^{-3}$ | 1.00 |
| 15 | 910 | 900.00 | 10.00 | 1.01 |
| 16 | 850 | 867.50 | 17.50 | 1.02 |
| 17 | 761 | 761.00 | $< 10^{-3}$ | 1.00 |
| 18 | 844 | 875.00 | 19.00 | 1.02 |
| 19 | 412 | 420.74 | 8.736 | 1.02 |
| 20 | 750 | 755.00 | 5.000 | 1.01 |
| **avg.** |  |  | 7.26 | 1.01 |
| **max.** |  |  | 19.00 | 1.04 |
| **min.** | | | $< 10^{-3}$ | 1.00 |

Figure 7e shows the reprojection error averaged across all frames for each of the 20 marmoset keypoints. The mean error for each keypoint ranged from 6.62 to 14.63 pixels, and the grand mean across all keypoints was 11.47 pixels. The nose and both eyes showed relatively low errors (6.62–7.08 pixels), whereas the right ear and right wrist (keypoint 11) showed the largest errors (14.58 and 14.63 pixels, respectively). The 95th percentile values ranged from 15.41 to 31.68 pixels, with the largest value observed for the left ankle (31.68 pixels).

### Geometric accuracy of the 3D environment model

To evaluate the dimensional accuracy of the created 3D environment model, dimensions of each part of the 3D model were compared with those in the blueprints of the experimental environment (Table 3). Among the 19 measurement points, 9 matched the blueprint values (absolute error *<* 10^−3^ mm), whereas the remaining 10 showed a mean error of approximately 13.3 mm and a maximum error of 19.0 mm. The dimensional ratios (3D model/blueprint) were within the range of 1.00–1.04 for all locations, and all parts deviated slightly in the direction of being larger than the blueprint dimensions. The maximum dimensional ratio of 1.04 occurred for the part with a blueprint value of 405 mm (No. 11), for which the absolute error was 15.74 mm. The mean absolute error across all measurement points was 7.26 mm (mean dimensional ratio, 1.01), indicating that the 3D environment model reproduced the blueprint dimensions with an accuracy of approximately 1%.

### Validation of consistency in experimental-environment reconstruction

The preceding sections evaluated each component of the dataset individually: the 3D marmoset model, 3D keypoint estimation, and 3D environment model. In this section, we evaluate the overall reproducibility of the integrated digital-twin environment. We compared real recorded images with rendered images in which the marmoset was placed in the 3D environment according to the estimated pose, and calculated SSIM and PSNR.

To generate computer-rendered images corresponding to the real video frames (the left panel of Figure 8a), we reproduced the eight environment cameras used in the experimental setup, as shown in the middle panel of Figure 8a. Because the marmoset was visible in the real video frames, the pose of the 3D marmoset model was also set using the estimated 3D keypoints. The resulting rendered images therefore corresponded to the real frames in terms of both camera viewpoint and marmoset pose. An example output used to calculate SSIM and PSNR is shown in the right panel of Figure 8a. The image mainly shows the SSIM map, in which white regions indicate structurally similar regions with SSIM scores close to 1, and black regions indicate dissimilar regions with scores close to 0. For this image pair, the SSIM was 0.596 and the PSNR was 18.48 dB.

**Figure 8.**
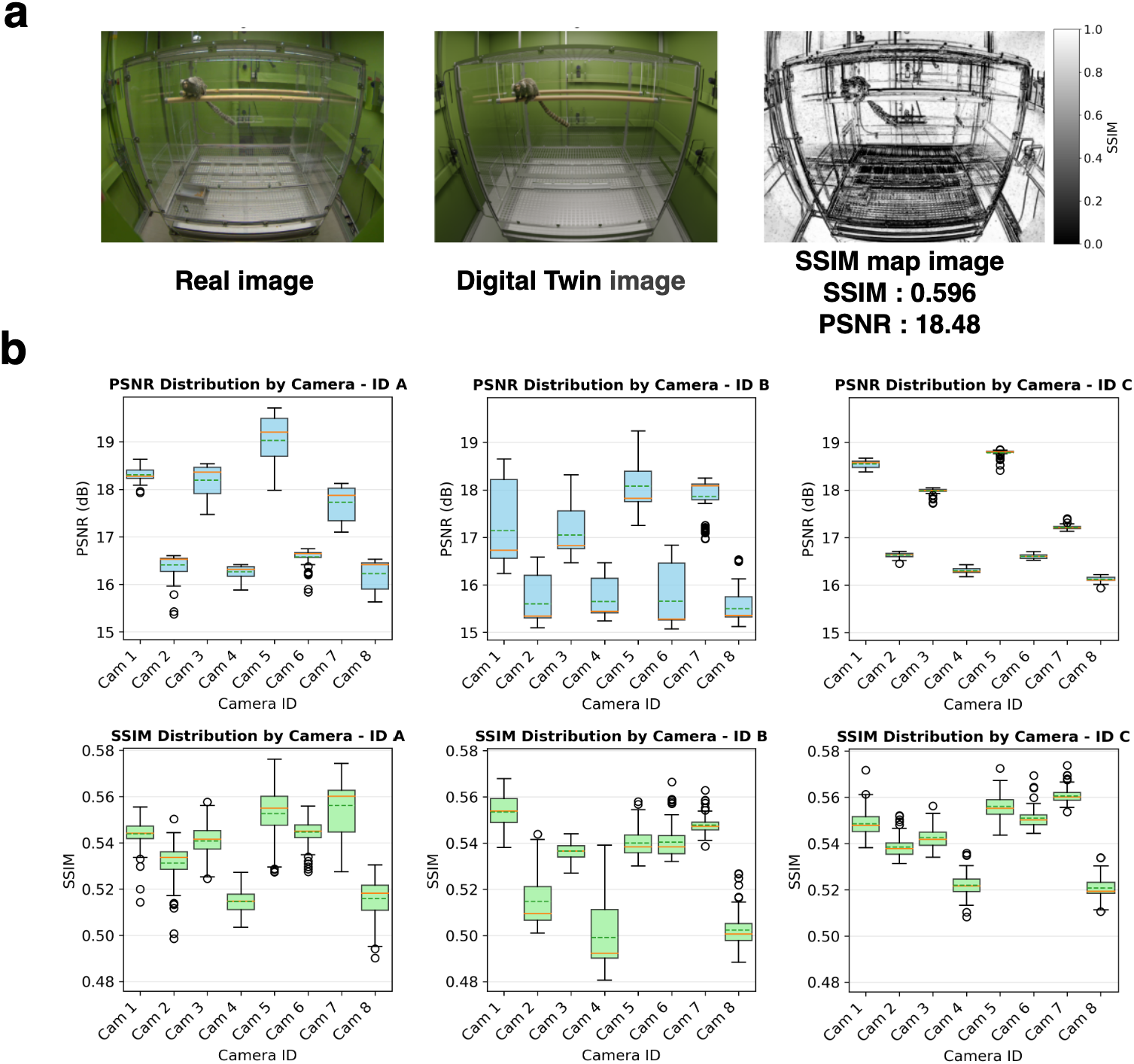
Validation of consistency in experimental-environment reconstruction. (a) Example used to calculate SSIM and PSNR. The left image shows a photograph of the marmoset in the experimental environment, the middle image shows a rendered image of the 3D marmoset created in this study, and the right image shows the calculated SSIM map. The SSIM and PSNR scores are shown at the top of the image. For this example image pair, the SSIM was 0.5962 and the PSNR was 18.48 dB. (b) Results of calculating PSNR and SSIM from image pairs such as those shown in panel a, using 100 frames for each image set. The upper row shows PSNR, and the lower row shows SSIM as box plots. The horizontal axis shows each camera, and the vertical axis shows the PSNR or SSIM score. Individuals A, B, and C are arranged from left to right. The green dashed line indicates the mean, and the orange solid line indicates the median.

Figure 8b shows the distributions of SSIM and PSNR values calculated from 100 randomly sampled frames for each combination of the eight cameras and three individuals. The upper row shows PSNR, and the lower row shows SSIM. For each individual, values are shown as box plots for each camera viewpoint, with individuals arranged from left to right as A, B, and C. Across all samples, the mean SSIM was approximately 0.536 and the mean PSNR was approximately 17.06 dB. The distributions of both metrics were broadly similar across the three individuals, with no substantial differences in SSIM or PSNR observed among individuals. These results show that the rendered images captured the overall scene structure but did not fully reproduce the pixel-level appearance of the real images.

## Usage Notes

The resources described in this Data Descriptor have two current limitations in their scope of application. First, the CT data underlying the 3D body model were obtained from one individual, whereas the behavioral videos were recorded from three different individuals. The resource should therefore be interpreted not as a strictly individual-specific digital twin, but as a high-fidelity 3D platform that can be synchronized with behavioral data. Second, the current pose synchronization and technical validation primarily target conditions in which a single individual is present in the cage. This is because 2D estimation, particularly for tail keypoints, can become unstable when multiple individuals are close together and fall within the same bounding box. Even within this scope, however, the ability to stably integrate the body surface, fur, rig, and spatial relationship with the environment is important for high-resolution visualization of marmoset body–environment interactions and provides a practical foundation for future extensions to multi-animal settings.

## Data availability

The dataset will be made available from Zenodo upon acceptance of this article and is currently available from an AWS S3 bucket at https://digitaltwin-marmoset.s3.ap-northeast-1.amazonaws.com/index.html. The Zenodo archive contains representative files and metadata, whereas the full dataset, including the complete video data, is hosted on AWS because of its large size. The total size of the full dataset hosted on AWS is approximately 920 GB. All files are released under CC-BY. Data and related resources for this study are available through the project website at https://nakaelab.github.io/marmoset-digital-twin/.

## Code availability

The 2D and 3D keypoint estimation pipeline, Python scripts for measuring the dimensions of the 3D environment model, and scripts for reproducing 3D poses in Blender are available in the GitHub repository 3D_digital_twin_marmoset_code (https://github.com/iwtkk/3D_digital_twin_marmoset_code.git). OpenCV, which was used for camera calibration, and Anipose, which was used for spatiotemporal smoothing of 3D keypoints, are used in the code in the GitHub repository. The following software was used in this study: Blender v4.5.0 for surface-model creation, rigging, environment-model construction, and rendering; MMDetection v2.26.0 for training the bounding-box detector; MMPose v0.29.0 for training the HRNet-W48-DEKR 2D pose estimation model; MATLAB R2023b for CT-segmentation preprocessing; and 3D Slicer 4.11 with the SlicerRT toolkit plugin for calculating the Dice coefficient and Hausdorff distance.

## Acknowledgements

We gratefully acknowledge the central institute for experimental medicine and life science and Dr. Takashi Inoue for providing the marmoset samples. We thank Dr. Teppei Ebina and Dr. Reona Kobayashi for valuable discussions and constructive feedback on this study. We also thank MELTA Inc. for their support in the 3D modeling of the marmoset. This work was supported by JST FOREST Program under Grant Number JPMJFR241Q to TK; JSPS KAKENHI under Grant Numbers JP22H05163 and JP24K03256 to K.N.; AMED under Grant Numbers JP22zf0127007, JP24wm0625414, and JP24wm0625404 to K.N.

## Author information

### Contributions

K.I. was responsible for data curation, formal analysis, investigation, methodology, resources, software, validation, visualization, and writing–review & editing. T.K. was responsible for data curation, formal analysis, investigation, resources, and software. C.C.C. was responsible for data curation and resources. D.T. was responsible for formal analysis, methodology, and software. A.N. was responsible for data curation and resources. T.M.N. was responsible for data curation and resources. J.H. was responsible for data curation and resources. K.N. contributed to conceptualization, formal analysis, funding acquisition, investigation, methodology, project administration, resources, software, supervision, validation, and writing–review & editing.

